# Proprioceptive and visual motion detection acuity contribute to children’s dynamic postural control

**DOI:** 10.1101/2025.04.29.651175

**Authors:** Antonella Iannotta, Scott J. Mongold, Esranur Yildiran Carlak, Christian Georgiev, Pierre Cabaraux, Dorine Van Dyck, Gilles Naeije, Marc Vander Ghinst, Jennifer Foucart, Nicolas Deconinck, Mathieu Bourguignon

## Abstract

Acquiring efficient postural control strategies is key to children’s proper motor development. For that, the brain needs to continuously integrate sensory information and convert it into corrective motor commands. Although this entire process naturally hinges on the reliability of early senses, very few studies have investigated early sensory acuity and its role in postural stability during development. Clarifying this could lead to a better understanding of conditions, such as developmental coordination disorder (DCD), where the impairment of balance control is substantial.

Here, we tested 25 typically developed school-aged children with a Visual Motion Detection test (VMDT), an ankle Joint Position Sense test (aJPST), force-plate assessed posturography, and the Movement Assessment Battery for Children - Second edition (MABC-2).

We found a significant correlation between the balance score of the MABC-2 and both VMDT score (*r* = 0.60, *p* = 0.003) and aJPST score (*r* = -0.47, *p* = 0.02). However, no such relationship was found between the force-plate assessed sway amplitude during upright standing and the two sensory acuity scores. Importantly, the MABC-2 balance scores were associated with upright stability, but only to a limited extent.

Given that the MABC-2 balance component factors in static and dynamic balance while posturography focuses only on static balance, our results point at a key role of early sensory acuity for dynamic balance. Together, these findings bring attention to possible clinical tools for motor impairment detection and subsequent rehabilitation strategies during development.

## 1. INTRODUCTION

Postural stability refers to the ability to maintain a specific position of the body and achieve balance through coordinated actions, and it relies on three sensory systems: the visual, proprioceptive and vestibular systems. These three fundamental perceptual systems generate information that enables the localization of the body in space and more specifically, the position of each body segment [1], enabling the maintenance of the center of gravity within the body’s base of support. The efficiency of each of these three systems and their integration varies across individuals and according to different circumstances such as changes in environmental constraints (e.g., standing in darkness decreases visual weight) or even by top-down attention [2].

As in adults, postural control in children is the result of a complex process of integration of sensory information from these three main sensory systems [3]. However, children do not have the same postural responses as adults until middle childhood (6-11 years) or later [4]. Sensory systems mature at different rates, and each child preferentially uses information from one or more systems [5]. For instance, children between the ages of 4 and 6 rely predominantly on visual input for maintaining a bipedal stance [6]. It is only between 7 and 10 years of age that vestibular and proprioceptive inputs are integrated in a manner comparable to adults [7]. The critical roles of visual acuity and proprioception in coordinated movement and postural stability are well-documented [8]. However, the relationship between early sensory acuity and postural stability remains underexplored, leaving gaps in understanding how sensory deficits affect balance maintenance in children.

Considering the above, the present study aims to evaluate to what extent postural stability hinges on early sensory acuity in typically developing children. Specifically, we evaluated typically developed school-aged children with a Visual Motion Detection test (VMDT), an ankle Joint Position Sense test (aJPST), posturography, and the Movement Assessment Battery for Children - Second edition (MABC-2; ref). The aJPST was performed at the ankle joint given its crucial role in balance maintenance [7], [9]. Likewise, the VMDT was selected given the relevance of visual motion detection acuity for upright balance maintenance [10], [11]. We hypothesized that poor visual motion detection acuity and ankle proprioceptive acuity would be associated with poor motor performances, especially in terms of postural stability. In addition, we hypothesized that the amplitude of postural sways obtained by posturography would be predictive of the MABC-2 score for postural stability.

This research may guide future investigations into conditions marked by significant postural instability. A striking and understudied example of this is Developmental Coordination Disorder (DCD). This neurodevelopmental disorder affects around 5-6% of children worldwide and is characterized by poor motor abilities and difficulty learning new motor skills, including poor static and dynamic balance and poor coordination [12]. Moreover, children with DCD show weaker visual reweighting, no advanced multisensory integration and delayed responses to multisensory stimuli compared with typically developing children [13].

By highlighting the fundamental role of early sensory acuity in postural control, our findings may pave the way for targeted interventions that address sensory deficits to improve motor outcomes in children, particularly those with conditions such as DCD.

## 2. MATERIALS AND METHODS

### 2.1 Participants selection and study approval

Twenty-five typically developing children (mean ± SD age, 8.3 ± 2.3 years, range 5-12 years, 12 females) were recruited to participate in the study. They had no history of movement disorders and were generally healthy as reported by their parents or legal guardian. The following exclusion criteria were employed: balance disorders, psychiatric disorders, and musculoskeletal injuries. The study was approved by the *ULB-Hôpital Erasme Ethics Committee* (P2023/313, CCB B4062023000180). Each child’s legal representative gave written informed consent before participation in accordance with the Declaration of Helsinki. The measurements were carried out at the ULB Hôpital Erasme, Brussels, Belgium. Participants received a gift card as compensation for their participation.

### 2.2 Experimental protocol

Subjects underwent a VMDT, an aJPST and an evaluation of the bipedal stance through posturography. In addition, the MABC-2 [14] was used to assess motor skills.

For the VMDT, participants sat in front of a 17-inch computer screen, with their eyes ∼90 cm away from the screen. The VMDT assessed their ability to recognize the vertical movement (either upwards or downwards) of a black–white Gabor patch oriented horizontally (full width at half maximum: 5.06 cm, corresponding to a visual angle of 3.22 degrees, spatial frequency: 1.16 cm^-1^ corresponding to 1.83 per degree of visual angle) and presented atop a gray background. In implementing the vertical movement, only the sine carrier was moved within a fixed Gaussian patch. The test included a total of 80 trials.

**Figure 1A** illustrates the VMDT task sequence. Each trial started with a gray screen for 0.5 s, followed by the presentation of the moving Gabor patch for 2 s. Then the participant was prompted to record the perceived motion direction via mouse click. The next trial started after the participant’s response. Shift period, defined as the time it takes for a stripe to move by a spatial period, was varied following a two-down/one-up staircase procedure. The initial shift period was 1 s, which a successful trial led to multiply by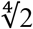 and a failed trial to divide by √2. As a result, the shift period converged onto a threshold that reflected each participant’s visual motion detection acuity.

**Figure 1.**
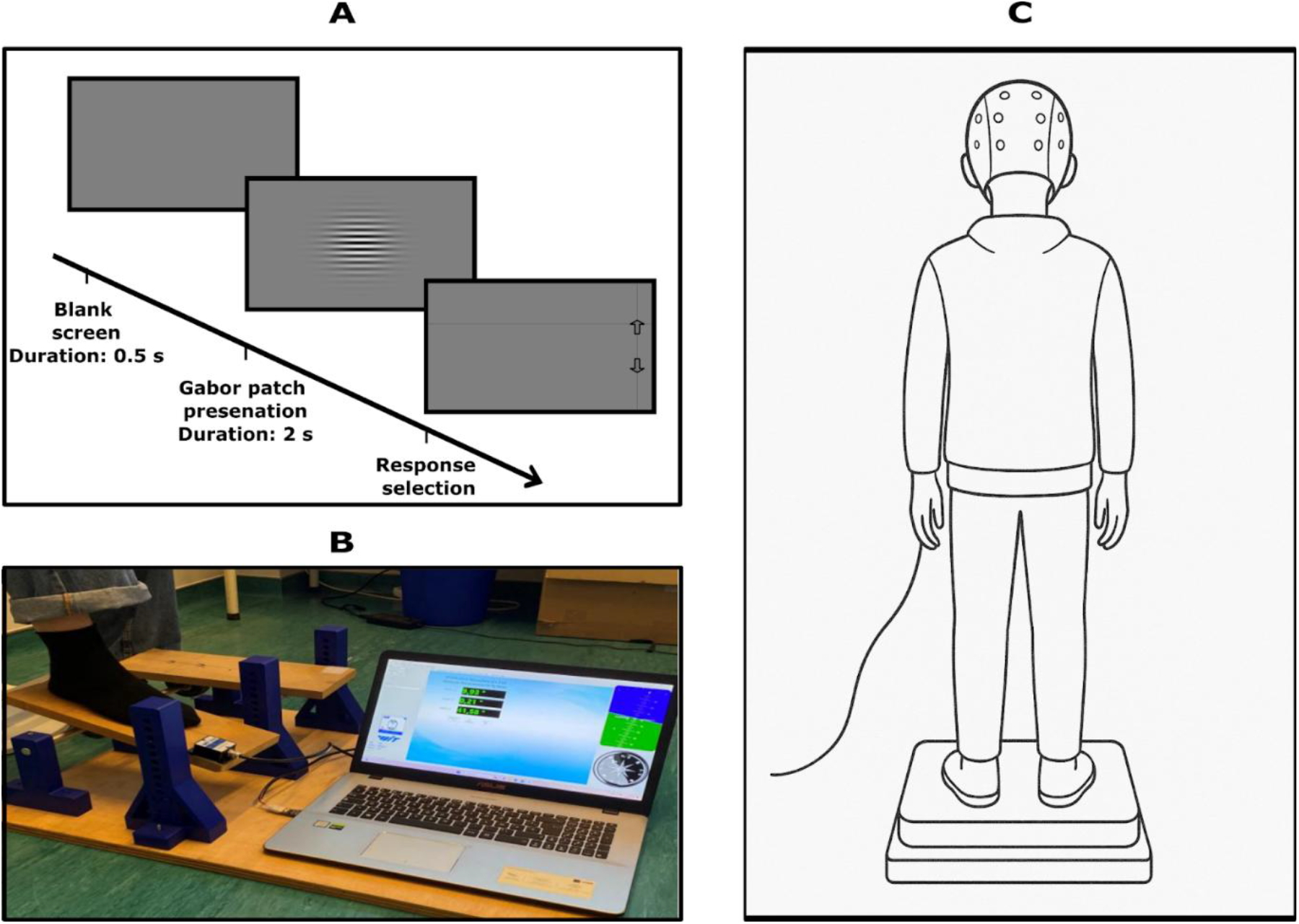
**A**. Time-course of the VMDT task. Each of the 80 trials started with a gray screen, followed by the visual stimulus, and then by the response selection screen. The Gabor patch moved in either the upwards or downwards direction. The participant moved the cursor to indicate the perceived direction of movement (adapted from [15]). **B**. aJPST set-up. Participants’ dominant foot rested on a movable plank attached to a custom-made structure that allowed rotation of the plank in the plantar/dorsiflexion directions. A gyroscope attached to the movable plank continuously monitored the angle. **C**. Posturography set-up. Participants were equipped with a 64 channel EEG-cap and stood on a force plate across four experimental conditions. EEG data was not analyzed for this study.

**Figure 1B** illustrates the setup for the aJPST. The test was conducted on the dominant leg, as determined by the subject’s self-report regarding the leg typically employed for kicking a ball. Participants sat on a chair with their legs at a 90° angle to the thighs. Participants were blindfolded using a mask. The tested foot was placed on a custom-made device consisting of a foot-sized plank that could rotate about an axis situated ∼6 cm underneath the ankle. A gyroscope (Wit motion HWT901B-RS485), featuring a precision of 0.05°, was fixed to a corner of the plank and used to record its orientation at 100 Hz during the whole duration of the procedure. Each trial consisted of three stages and began and ended with the ankle at the ‘neutral’ position, corresponding to the ankle oriented 90° relative to the shank. First, the ankle was passively mobilized to a predefined angle and held for ∼3 s. Second, the ankle was passively repositioned to the starting position for ∼ 3 s. Finally, the participant was asked to actively move their ankle to the previous position as accurately as possible, and to hold this position for ∼ 3 s. Passive mobilizations were performed at constant speed. Nine trials were performed in random order, at three different angles (dorsiflexion: 10° and 15°; plantar flexion: 10°).

**Figure 1C** illustrates the posturography setup. Participants stood atop a force plate (AccuSway-O, AMTI, Watertown, MA, USA) and underwent four different balance conditions: with the eyes open on a hard surface, with the eyes closed on a hard surface, with the eyes open on foam pads (Domyos, Decathlon, Villeneuve-d’Ascq, France) and with the eyes closed on foam pads. Participants were asked to maintain their balance and stay relaxed. Each participant performed 8 randomized trials (each condition twice) of 5 minutes each, for a total of 40 minutes of recording. During each trial, the force plate measured the forces and force moments applied at ground level at a sampling frequency of 1000 Hz.

The *Movement Assessment Battery for Children – Second Edition* (MABC-2) [14] assesses three major domains: manual dexterity, static and dynamic balance and the capacity of aiming and catching. Participants completed a total of eight subtests, according to their age group. Each participant was given a training session to familiarize themselves with the task.

### 2.3 Data processing

Unless specified otherwise, the data analysis was performed using custom scripts in Matlab (version R2024a; Mathworks, Natick, MA, USA).

#### VMDT

The VMDT score was calculated using the geometric mean of the shift period in the last 30 trials. Participants with higher VMDT score are deemed to possess a better visual motion detection acuity compared to participants with lower scores.

#### aJPST

The aJPST score was calculated as the mean of the relative angle reproduction error across all trials. The relative angle reproduction error was itself the absolute value of the difference between the imposed and reproduced angles divided by the imposed angle. Increased aJPST scores indicate larger errors and thus correspond to worse proprioceptive acuity.

#### Posturography

The position of the center of pressure (CoP) was calculated using force plate data. CoP time-series were filtered between 0.2 and 10 Hz using a digital filter. To ensure that only data from the bipedal stance were processed, recordings were taken 10 seconds after ascent and 10 seconds before descent from the force plate. To quantify postural instability, we assessed the standard deviation of the CoP along the anterior-posterior axis (sdCoP_AP_) for each recording separately. Higher sdCoP_AP_ indicated greater postural instability. Asingle value per condition was obtained as the mean of sdCoP_AP_ across the two trials of that condition.

#### MABC-2

Motor performance data were obtained from the raw scores of the MABC-2 tests. These scores were converted to standard scores according to the age of the participants, using the conversion tables in the MABC-2 manual. The standard scores were aggregated within the same components assessing motor performance. Each component was converted into standard scores and percentiles using another conversion table, providing data on static and dynamic balance, aiming and catching ability, and manual dexterity. Finally, the sum of the standard scores was converted to percentiles using a final conversion table, providing an overall measure of each participant’s motor performance. These standard scores indicate an individual’s relative position within a reference distribution.

### 2.4 Statistical analyses

Statistical analyses were performed using RStudio (version 2024.04.2+764). Prior to statistical modeling, VMDT, aJPST and sdCoP_AP_ scores were log-transformed to better approximate a normal distribution, and to comply with the homoscedasticity assumption [15]. Scores were then corrected for outliers by setting values greater than 2.5 SD above or below the mean to this threshold.

Pearson’s correlations were used to assess the dependence of VMD acuity and proprioceptive error on the results of each section of the MABC-2.

We performed a linear mixed model analysis with *lme4* to evaluate how sdCoP_AP_ depends on five fixed effects: condition, age, VMDT score, aJPST score and MABC-2 balance score. We started with a null model that included only a different random intercept for each subject. The model was iteratively incremented with fixed effects and compared to the non-incremented model using a *χ*^2^ test. Effects were tested in a planned order: (1) condition, (2) age and, (3) its interaction with condition, (4) VMDT score and, (5) its interaction with condition, (6) aJPST score and, (7) its interaction with condition, (8) MABC-2 balance score, and (9) its interaction with condition. At every step, the added fixed effect was retained in the model if deemed significant (*p* < 0.05). Post-hoc comparisons for significant effects were conducted with t-tests for categorical fixed effects (condition) and Pearson’s correlations for continuous ones.

## 3. RESULTS

### 3.1 Relationship between motor abilities and sensory acuity

**Figure 2** presents the distribution of motor performance data from the MABC-2 expressed as a percentile of the scores in a reference population. Scores were 41 ± 28 % (manual dexterity; mean ± SD), 47 ± 27 % (aiming and catching), 61 ± 22 % (balance) and 48 ± 25 % (total score). The substantial standard deviation observed in the results could indicate that, even in the context of testing typically developed children, some may potentially be at risk of motor impairments, as indicated by the standardized score of the MABC-2. Two participants in the present study had a total score in the percentile 16, which is considered to be the upper limit of the “orange zone” [15]. This zone is characterized as the area in which children are at risk of motor impairments. Furthermore, both participants received a low score in one of the three sections (manual dexterity for one participant and aiming and catching for the other). With the exception of the aforementioned subjects, the remaining participants demonstrated a percentile score that did not indicate any risk of motor impairment.

**Figure 2.**
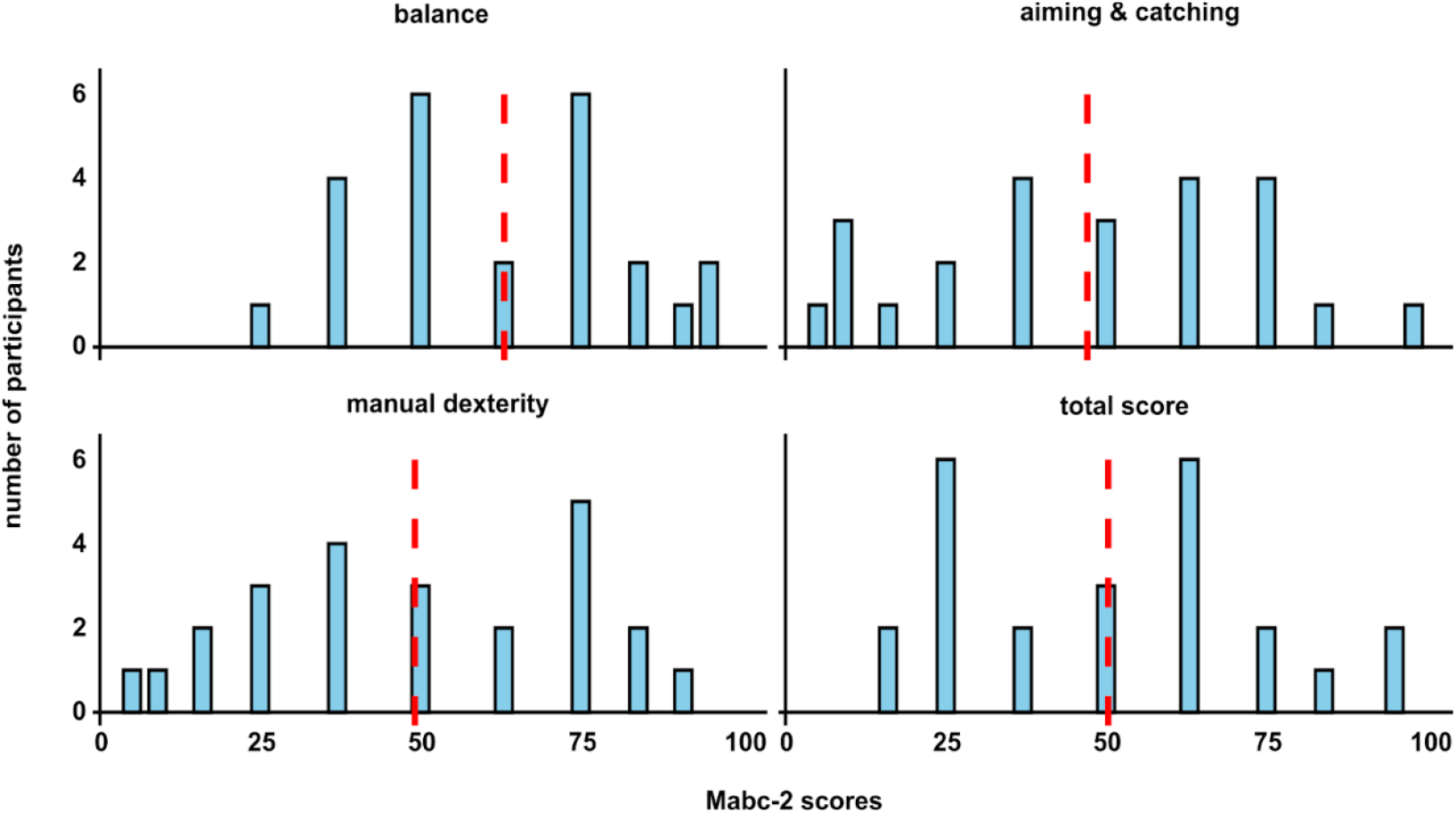
Distribution of the MABC-2 scores, expressed in percentile, across all participants. The red line in each graph represents the mean for each section.

Sensory acuity varied considerably between children, with VMDT scores characterized by a coefficient of variation of 76.8 % (mean ± SD shift period, 32.5 ± 24.9 s) and aJPST score by a coefficient of variation of 64.5 % (mean ± SD relative error, 52.2 ± 33.7 %).

**Figure 3** presents the results of the correlations between, on the one hand, VMDT and aJPST scores, and, on the other hand, the results of each section of the MABC-2. VMDT score correlated significantly with the balance score (*r* = 0.60, *p* = 0.003) and the total score (*r* = 0.49, *p* = 0.018). aJPST score correlated significantly with the balance score (*r* = -0.47, *p* = 0.02).

**Figure 3.**
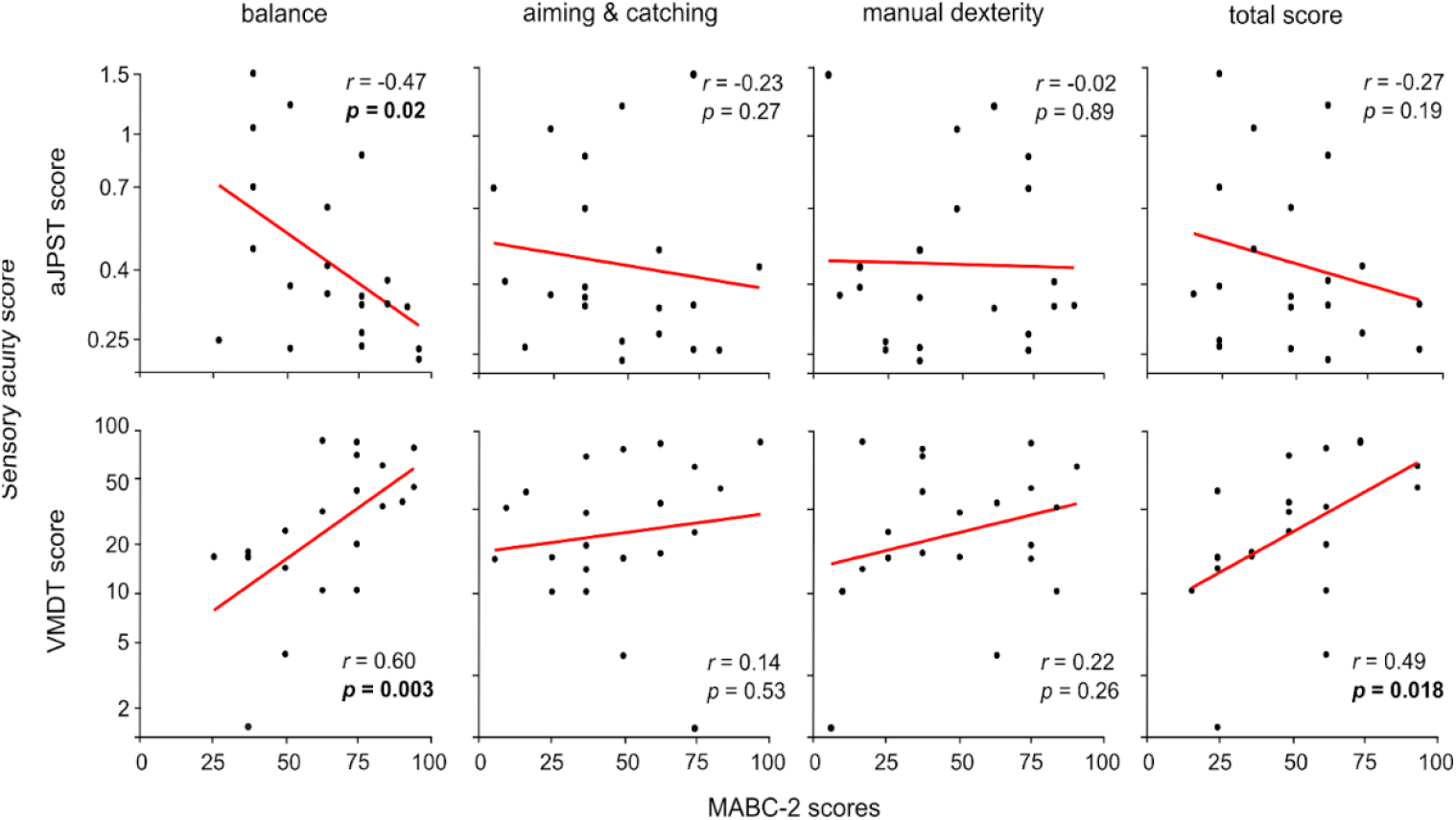
Relationship between sensory acuity (VMDT and aJPST scores) and MABC-2 scores. Black dots indicate individual participant’s values and the regression line across all participants is in red. Correlation value and the corresponding significance level for each association is indicated in the upper or bottom right corner.

### 3.2 Instability assessed with posturography

**Figure 4** presents the distribution of sdCoP_AP_ across participants in the four balance conditions. The linear mixed model revealed a significant effect of condition on sdCoP_AP_ (*χ*(3)^2^ = 87.1, *p* < 0.0001). The result of post-hoc comparisons between conditions is presented in **Figure 4**. As expected, these comparisons revealed that sdCoP_AP_ increased with condition complexity.

**Figure 4.**
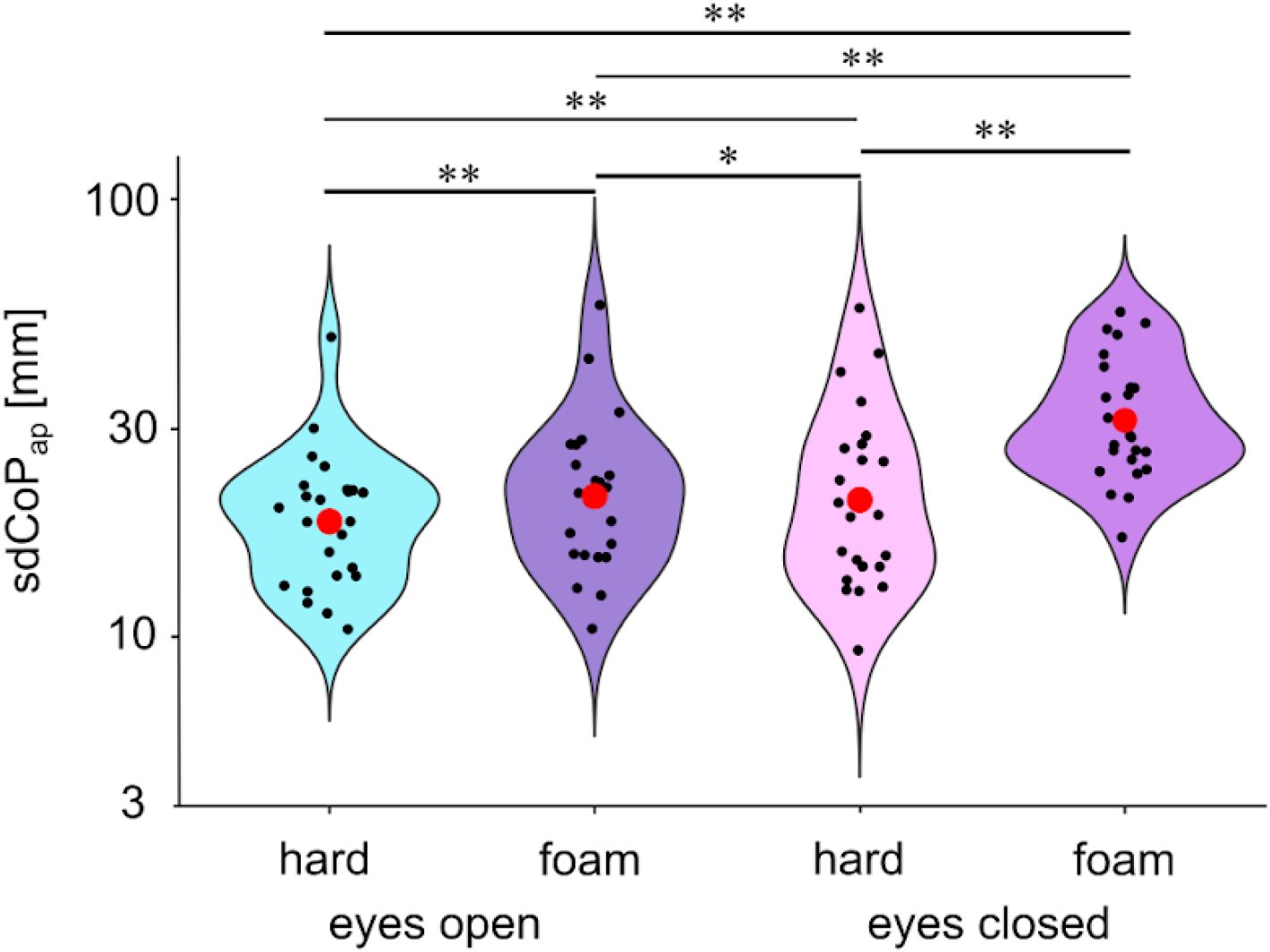
Distribution of sdCoP_AP_ in the four different conditions on a logarithmic scale. Children showed higher instability in the hardest condition (eyes-closed foam). Black dots indicate individual participants’ values and red dots the mean within conditions. Asterisks represent significance levels with: *, p < 0.05 and *^*^, p < 0.01

The linear mixed model revealed a significant effect of age on sdCoP_AP_ (*χ*(1)^2^ = 4.96, *p* = 0.02), which was not modulated by the standing condition (*χ*(3)^2^ = 5.15, *p* = 0.16). With increasing age, sdCoP_AP_ decreased in all conditions, indicating improved stability (see **Figure 5**).

**Figure 5.**
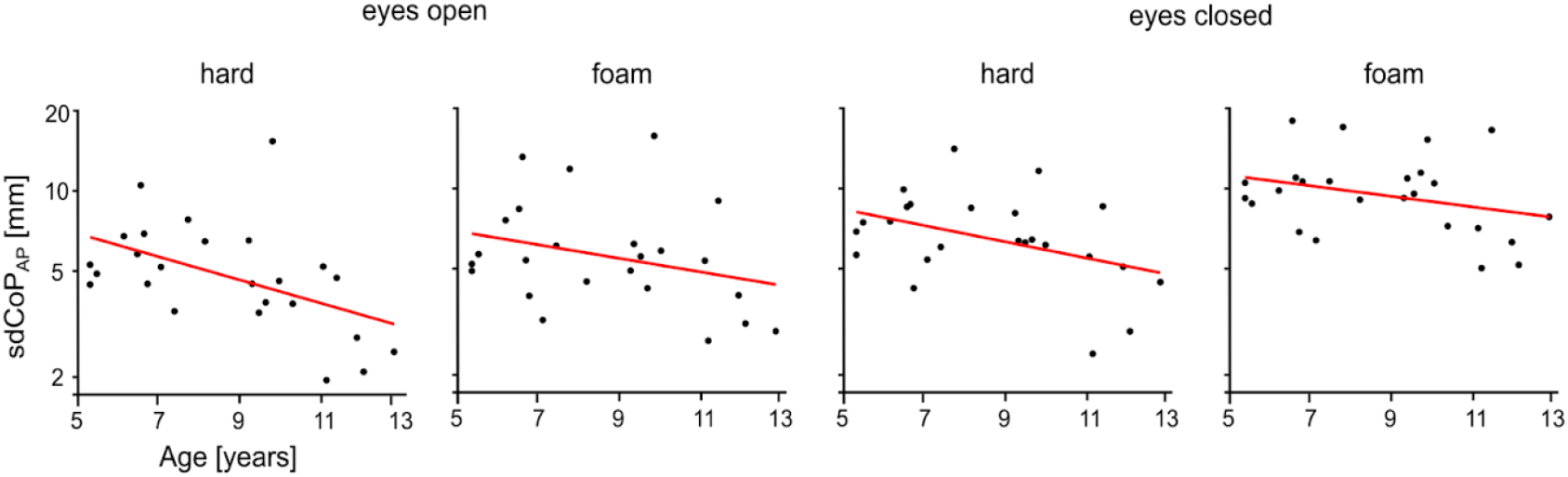
Relationship between sdCoP_AP_ and age in the four different conditions. Black dots indicate individual participants’ values and the regression line across all participants is in sred.

### 3.3 Relationship between instability and sensory acuity

The linear mixed model analysis assessing sdCoP_AP_ did not identify a significant effect of sensory acuity (VMDT score, *χ*(1)^2^ = 2.09, *p* = 0.14; aJPST score, *χ*(1)^2^ = 1.03, *p* = 0.30), nor a significant interaction thereof with the condition (VMDT score, *χ*(4)^2^ = 7.88, *p* = 0.09; aJPST score, *χ*(4)^2^ = 5.89, *p* = 0.20).

### 3.4 Relationship between instability and MABC-2

The linear mixed model analysis assessing sdCoP_AP_ identified a significant effect of the MABC-2 balance score (*χ*(1)^2^ = 3.67, *p* = 0.05), and no significant interaction thereof with the condition (*χ*(4)^2^ = 7.85, *p* = 0.09).

**Figure 6.** presents the correlation between the MABC-2 balance score and sdCoP_AP_. The correlation was negative in all conditions, albeit significant only in the eyes-closed hard condition (*r* = -0.42; *p* = 0.04).

## 4. DISCUSSION

In this study, we examined the relationship between sensory acuity and balance control, assessed with the MABC-2 and force plate-based posturography. We found that visual motion detection acuity and ankle proprioceptive acuity were significantly associated with the balance score of the MABC-2 (VMDT *p* = 0.003; aJPST *p* = 0.02), but not with force plate-assessed sway amplitude (VMDT *p* = 0.14, aJPST *p* = 0.30). Furthermore, the latter two measures of balance showed only a mild degree of association. Finally, confirmatory in nature, postural stability assessed with sway amplitude decreased with condition complexity, but improved with age within the 5-12 year age range.

**Figure 6.**
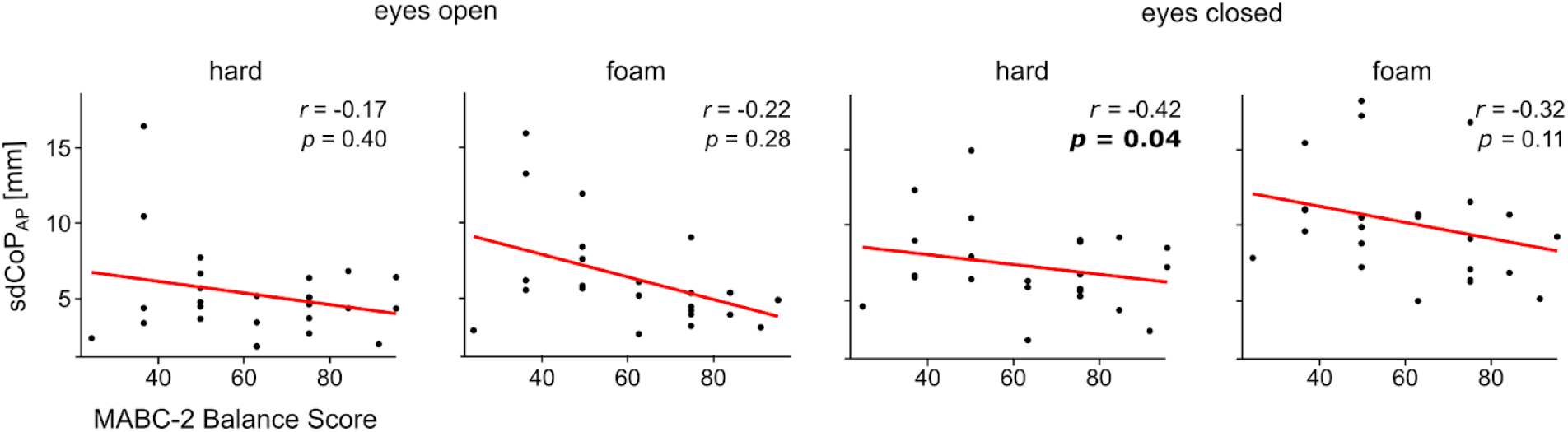
Relationship between sdCoP_AP_ and the balance score of the MABC-2 in the four different conditions. Graph layout as in Figure 5. Correlation value and corresponding significance level for each association is indicated in the upper right corner.

### 4.1 Importance of sensory acuity for motor performance

Although the present study was conducted in a cohort of typically developing children, notable variability in motor performance was observed, suggesting that even within a non-clinical population, subtle motor difficulties may be present.

A critical finding is the role of early sensory acuity in maintaining balance. That is, children with a higher score on the MABC-2 balance component displayed higher visual motion detection acuity and higher ankle proprioceptive acuity (i.e., lower errors on the aJPST). The association with visual motion detection acuity is in line with findings that children with DCD often present with visual deficits such as difficulties with fixation, abnormal eye movements, and binocular vision issues [16]. In the same vein, *Cheng* and colleague [17] reported that in children with DCD, poor visual performance has a negative impact on MABC-2 motor performance and daily life activities. As well, the association of the MABC-2 balance score with ankle proprioceptive acuity is in line with the fundamental role of proprioception in motor control. Ankle proprioception was demonstrated to contribute to balance, with extensive benefits of higher acuity, as has been reported in the context of sport performance and limitation of risks of injury [18] as well as for the limitation of balance impairment following stroke [19].

Critically, our data show that even in typically developing children, proprioception acuity at the ankle joint contributes to postural stability. These results align with the presence of proprioceptive deficits in individuals with DCD [20], [21]. Altogether, our findings support the notion that effective balance control may depend on the reliability of both the visual and proprioceptive systems. Therefore, future studies should determine the added value of VMDT and aJPST scores in clinical evaluations as predictive indicators for the early identification of motor deficits.

No significant correlations emerged between our measures of early sensory acuity and the two MABC-2 components other than balance. Given that our sensory tests assessed aspects of perception that are most relevant for balance maintenance, and less so for manual dexterity and catching abilities, our results are not surprising. Proprioceptive acuity of upper limb joints and aspects of visual acuity subtending visuomotor coordination could be relevant for manual dexterity and catching ability. Accordingly, wrist proprioceptive acuity was found to be altered both in adults and children with DCD [22], [23].This alteration correlated with the manual dexterity component of the MABC-2 in children with DCD [22], and with levels of body coordination in adults, as assessed with the Bruininks-Oseretsky Test of Motor Proficiency [23]. Likewise, fine motor skills in healthy adults were reported to be associated with a combined measure of near visual acuity, near contrast sensitivity, and disability glare [24].

### 4.2 Difference between MABC-2 and force plate-derived posturography

Although there was a negative trend between sway amplitude and MABC-2 scores, the correlation between these two aspects was only significant in the eyes closed condition, with a modest magnitude. Two main factors could explain the weakness of this association: the limited reproducibility of the measures, and the difference in balance tasks used. The MABC-2 scores are subject to some degree of bias since the examiner can influence the child’s performance through disparities in instructional, motivational and rating styles [25]. Still, the test has been demonstrated to possess excellent test-retest reliability, with intraclass correlation coefficients (ICCs) ranging from 0.83 to 0.99 [26], [27]. In contrast, force plates provide fully objective measures of balance. Despite that fact, the reliability of force plate-based posturography is similar to that of the MABC-2. Indeed, ICCs of around 0.80–0.95 were reported for measures tightly related to sdCoP_AP_ and vCoP_AP_ assessed in stroke patients [28] and in older adults [29]. Accordingly, the similarity in ICCs reported for MABC-2 balance scores and force plate-derived measures suggests that the limited association between the two is explained by other sources of difference. The explanation could be the difference in the balance tasks for the two tests, where posturography focuses solely on static balance while the MABC-2 integrates more dynamic tasks. In line with this idea, Liu et al. [30] found weak correlations between static and dynamic balance skills in a cohort of 4 to 5 years old children. The same conclusion was reached by a meta-analysis conducted over the lifespan [31].

### 4.3 Differential role of sensory acuity for static and dynamic balance

In our data, early sensory acuity was predictive of stability assessed with the MABC-2 but not with posturography. In light of the considerations developed above, this suggests that such acuity is more important for dynamic than static tasks. This distinction aligns with the broader understanding that dynamic balance relies heavily on feedback control [32]. In such tasks, the nervous system uses continuous sensory input to update internal models and generate timely motor corrections. As a result, accurate sensory information is essential for effective dynamic balance control. In contrast, static balance typically requires less real-time sensory feedback and receives a contribution from feedforward control strategy [33].

The difference in sensory demands between dynamic and static balance is also reflected in previous findings on DCD. Indeed, dynamic postural control was shown to be a great challenge for children suffering from DCD [34], whereas under normal conditions, static balance control is not [35]. Only under difficult, unattended, or novel situations do such children seem to suffer from increased postural sway [36]. Although we did not include children with DCD, our findings may provide some important elements of information for existing theories of this disorder. One influential hypothesis attributes DCD to deficits in internal models [37]. However, given the link between sensory acuity and dynamic balance, our data suggests that impaired sensory acuity may be a more fundamental issue. Such a deficit could compromise the development of accurate internal models, thereby contributing to the motor difficulties observed in DCD.

Finally, another potential explanation for the lack of significant relationship between sensory acuity and static balance stability may lie in the postural control strategy of the central nervous system. Although it would appear natural that sway amplitude be minimized, some models of feedback control suggest that, instead, humans attempt to minimize muscle activity [32], [38] with feedback control operating in an intermittent manner [39], [40]. Accordingly, sway amplitude may reflect an accepted level of instability rather than the minimal instability theoretically achievable through precise sensory feedback generated corrections.

### 4.4 Maturation of postural stability

Force plate posturography results showed that instability increases under more complex conditions,such as on unstable surfaces or when visual information is absent, and decreases as children age. The effect of condition complexity is a classic finding in the field and has been reported extensively in adults [15], [41], [42] and children [43], [44]. The maturation with age is also well documented [45], [46], especially between age 6 and 10 [47], with adult-like behavior appearing around age 7–8 [48], [49].

### 4.5 Limitations and perspectives

The main limitation of our study lies in that we did not control for the role of attention in our sensory and balance assessments. SClearly, some children found it challenging to sustain concentration and complete each task block, which required ∼5 minutes of continuous attention. Future studies should consider shortening individual tasks and adapting them into more child-friendly implementations using colours/cartoons. Further research is required to elucidate the developmental trajectory of sensory-motor integration and explore the potential of sensory acuity measures as predictive markers for motor impairments, particularly in populations at risk for DCD. Notwithstanding the methodological limitations, these results lay the foundation for targeted intervention strategies and future longitudinal research. This can further refine diagnostic tools and therapies, thereby reinforcing the paradigm that robust sensory integration is the basis of effective motor function

### 4.6 Conclusions

In conclusion, our study suggests that both visual motion detection and proprioceptive acuity are associated with balance control, with age playing a significant role in this relationship. The results further highlight the key distinction between static and dynamic postural control, where only the later seems to benefit from early and better sensory acuity. Collectively, our results indicate that future studies should determine the added value of early sensory acuity assessments as clinical tools to benefit diagnosis and treatment of children with motor disorders.

## Acknowledgment

Scott Mongold was supported by an Aspirant Research Fellowship awarded by the F.R.S.-FNRS (Brussels, Belgium; grant FC 46249). Christian Georgiev was supported by an Aspirant Research Fellowship awarded by the Fonds de la Recherche Scientifique (F.R.S.- FNRS; Brussels, Belgium; grant 1.A.211.24F). Pierre Cabaraux was supported by a Clinical Researcher Fellowship awarded by the F.R.S.-FNRS (Brussels, Belgium; grant 40024164). Gilles Naeije is postdoctorate Clinical Master Specialist at the FRS-FNRS (Brussels, Belgium). The project was supported by grants of the Fonds de la Recherche Scientifique (F.R.S.-FNRS, Brussels, Belgium; grant MIS F.4504.21), and of the Brussels-Wallonia Federation (Collective Research Initiatives grant) awarded to Mathieu Bourguignon.

## REFERENCES

[1] “Horak, F. B., & Macpherson, J. M. (1996). Postural Orientation and Equilibrium. In L. B. Rowell, & J. T. Sheperd (Eds.), Handbook of Physiology, Section 12. Exercise: Regulation and Integration of Multiple Systems (pp. 255–292). New York: Oxford University Press.”

[2] “Peterka RJ. Sensory integration for human balance control. Handb Clin Neurol. 2018;159:27–42. doi: 10.1016/B978-0-444-63916-5.00002-1. PMID: 30482320.”

[3] T. C. K. Cheung and M. A. Schmuckler, “Multisensory and biomechanical influences on postural control in children,” J. Exp. Child Psychol., vol. 238, p. 105796, Feb. 2024, doi: 10.1016/j.jecp.2023.105796.

[4] “Pierret J, Beyaert C, Paysant J, Caudron S. How do children aged 6 to 11 stabilize themselves on an unstable sitting device? The progressive development of axial segment control. Hum Mov Sci. 2020 Jun;71:102624. doi: 10.1016/j.humov.2020.102624. Epub 2020 Apr 25. PMID: 32452427.”

[5] W.-N. Bair, T. Kiemel, J. J. Jeka, and J. E. Clark, “Development of multisensory reweighting for posture control in children,” Exp. Brain Res., vol. 183, no. 4, pp. 435–446, Oct. 2007, doi: 10.1007/s00221-007-1057-2.

[6] “Sparto PJ, Redfern MS, Jasko JG, Casselbrant ML, Mandel EM, Furman JM. The influence of dynamic visual cues for postural control in children aged 7–12 years. Exp Brain Res. 2006 Jan;168(4):505–16. doi: 10.1007/s00221-005-0109-8. Epub 2005 Sep 7. PMID: 16151780.”.

[7] “Shumway-Cook A, Woollacott MH. The growth of stability: postural control from a development perspective. J Mot Behav. 1985 Jun;17(2):131–47. doi: 10.1080/00222895.1985.10735341. PMID: 15140688.”.

[8] D. N. Lee and E. Aronson, “Visual proprioceptive control of standing in human infants,” Percept. Psychophys., vol. 15, no. 3, pp. 529–532, May 1974, doi: 10.3758/BF03199297.

[9] F. Chen, C. Pan, C. Chu, C. Tsai, and Y. Tseng, “Joint position sense of lower extremities is impaired and correlated with balance function in children with developmental coordination disorder,” J. Rehabil. Med., vol. 52, no. 8, p. jrm00088, 2020, doi: 10.2340/16501977-2720.

[10] “Paulus WM, Straube A, Brandt T. Visual stabilization of posture. Physiological stimulus characteristics and clinical aspects. Brain. 1984 Dec;107 (Pt 4):1143–63. doi: 10.1093/brain/107.4.1143. PMID: 6509312.”.

[11] “Wade MG, Jones G. The role of vision and spatial orientation in the maintenance of posture. Phys Ther. 1997 Jun;77(6):619–28. doi: 10.1093/ptj/77.6.619. PMID: 9184687.”.

[12] Castellucci G., Singla R., “Developmental Coordination Disorder (Dyspraxia),” StatPearls [Internet]. Treasure Island (FL), Feb. 24, 2024.

[13] W.-N. Bair, T. Kiemel, J. J. Jeka, and J. E. Clark, “Development of Multisensory Reweighting Is Impaired for Quiet Stance Control in Children with Developmental Coordination Disorder (DCD),” PLoS ONE, vol. 7, no. 7, p. e40932, Jul. 2012, doi: 10.1371/journal.pone.0040932.

[14] Henderson, S. E., Sugden, D. A., & Barnett, A. L.Ray-Subramanian, “Movement Assessment Battery for Children: 2nd Edition (MABC-2),” 2007.

[15] “Cabaraux P, Mongold S, Georgiev C, Yildiran Carlak E, Garbusinski J, Naeije G, Vander Ghinst M, Bourguignon M. The confusing role of visual motion detection acuity in postural stability in young and older adults. Gait Posture. 2025 Feb 27;119:63–69. doi: 10.1016/j.gaitpost.2025.02.027. Epub ahead of print. PMID: 40043516.”.

[16] “Elena, Pinero-Pinto & Romero-Galisteo, Rita-Pilar & Sánchez González, María Carmen & Prieto, Escobio & Luque-Moreno, Carlos & Palomo-Carrión, Rocío. (2022). Motor Skills and Visual Deficits in Developmental Coordination Disorder: A Narrative Review. Journal of Clinical Medicine. 11. 7447. 10.3390/jcm11247447.”.

[17] “Cheng CH, Ju YY, Chang HW, Chen CL, Pei YC, Tseng KC, Cheng HY. Motor impairments screened by the movement assessment battery for children-2 are related to the visual-perceptual deficits in children with developmental coordination disorder. Res Dev Disabil. 2014 Sep;35(9):2172–9. doi: 10.1016/j.ridd.2014.05.009. Epub 2014 Jun 7. PMID: 24915646.”.

[18] “Han J, Anson J, Waddington G, Adams R, Liu Y. The Role of Ankle Proprioception for Balance Control in relation to Sports Performance and Injury. Biomed Res Int. 2015;2015:842804. doi: 10.1155/2015/842804. Epub 2015 Oct 25. PMID: 26583139; PMCID: PMC4637080.”.

[19] “Cho JE, Kim H. Ankle Proprioception Deficit Is the Strongest Factor Predicting Balance Impairment in Patients With Chronic Stroke. Arch Rehabil Res Clin Transl. 2021 Nov 2;3(4):100165. doi: 10.1016/j.arrct.2021.100165. PMID: 34977547; PMCID: PMC8683870.”.

[20] “Li KY, Su WJ, Fu HW, Pickett KA. Kinesthetic deficit in children with developmental coordination disorder. Res Dev Disabil. 2015 Mar;38:125–33. doi: 10.1016/j.ridd.2014.12.013. Epub 2015 Jan 7. PMID: 25576876.”.

[21] “Tran HT, Li YC, Lin HY, Lee SD, Wang PJ. Sensory Processing Impairments in Children with Developmental Coordination Disorder. Children (Basel). 2022 Sep 22;9(10):1443. doi: 10.3390/children9101443. PMID: 36291382; PMCID: PMC9600147.”.

[22] “Tseng YT, Tsai CL, Chen FC, Konczak J. Wrist position sense acuity and its relation to motor dysfunction in children with developmental coordination disorder. Neurosci Lett. 2018 May 1;674:106–111. doi: 10.1016/j.neulet.2018.03.031. Epub 2018 Mar 17. PMID: 29559417.”.

[23] “Tseng YT, Lin YH, Chen YW, Tsai CL, Chen FC. Impaired wrist position sense is linked to motor abnormalities in young adults with a probable developmental coordination disorder. Neurosci Lett. 2022 Feb 16;772:136446. doi: 10.1016/j.neulet.2022.136446. Epub 2022 Jan 7. PMID: 34999167.”.

[24] “Granados-Delgado P, Casares-López M, Martino F, Anera RG, Castro-Torres JJ. The Role of Visual Performance in Fine Motor Skills. Life (Basel). 2024 Oct 23;14(11):1354. doi: 10.3390/life14111354. PMID: 39598153; PMCID: PMC11595507.”.

[25] “Hadwin KJ, Wood G, Payne S, Mackintosh C, Parr JVV. Strengths and weaknesses of the MABC-2 as a diagnostic tool for developmental coordination disorder: An online survey of occupational therapists and physiotherapists. PLoS One. 2023 Jun 2;18(6):e0286751. doi: 10.1371/journal.pone.0286751. PMID: 37267388; PMCID: PMC10237484.”

[26] “Wuang YP, Su JH, Su CY. Reliability and responsiveness of the Movement Assessment Battery for Children-Second Edition Test in children with developmental coordination disorder. Dev Med Child Neurol. 2012 Feb;54(2):160–5. doi: 10.1111/j.1469-8749.2011.04177.x. PMID: 22224668.”

[27] “Ghayour Najafabadi M, Saghaei B, Shariat A, Ingle L, Babazadeh-Zavieh SS, Shojaei M, Daneshfar A. Validity and reliability of the movement assessment battery second edition test in children with and without motor impairment: A prospective cohort study. Ann Med Surg (Lond). 2022 Apr 28;77:103672. doi: 10.1016/j.amsu.2022.103672. PMID: 35638021; PMCID: PMC9142614.”

[28] “Aryan R, Inness E, Patterson KK, Mochizuki G, Mansfield A. Reliability of force plate-based measures of standing balance in the sub-acute stage of post-stroke recovery. Heliyon. 2023 Oct 14;9(10):e21046. doi: 10.1016/j.heliyon.2023.e21046. PMID: 37886778; PMCID: PMC10597864.”

[29] “Li, Z., Liang, Y.-Y., Wang, L., Sheng, J., and Ma, S.-J. (2016). Reliability and Validity of Center of Pressure Measures for Balance Assessment in Older Adults. J. Phys. Ther. Sci. 28 (4), 1364–1367. doi:10.1589/jpts.28.1364.”

[30] “Liu R, Yang J, Xi F, Xu Z. Relationship between static and dynamic balance in 4-to-5- year-old preschoolers: a cross-sectional study. BMC Pediatr. 2024 May 9;24(1):295. doi: 10.1186/s12887-024-04747-6. PMID: 38724964; PMCID: PMC11080223.”

[31] “Kiss R, Schedler S, Muehlbauer T. Associations Between Types of Balance Performance in Healthy Individuals Across the Lifespan: A Systematic Review and Meta-Analysis. Front Physiol. 2018 Sep 28;9:1366. doi: 10.3389/fphys.2018.01366. PMID: 30323769; PMCID: PMC6172339.”.

[32] R. Chiba, K. Takakusaki, J. Ota, A. Yozu, and N. Haga, “Human upright posture control models based on multisensory inputs; in fast and slow dynamics,” Neurosci. Res., vol. 104, pp. 96–104, Mar. 2016, doi: 10.1016/j.neures.2015.12.002.

[33] “Gatev P, Thomas S, Kepple T, Hallett M. Feedforward ankle strategy of balance during quiet stance in adults. J Physiol. 1999 Feb 1;514 (Pt 3)(Pt 3):915–28. doi: 10.1111/j.1469-7793.1999.915ad.x. PMID: 9882761; PMCID: PMC2269093.”

[34] P. H. Wilson, S. Ruddock, B. Smits-Engelsman, H. Polatajko, and R. Blank, “Understanding performance deficits in developmental coordination disorder: a meta-analysis of recent research,” Dev. Med. Child Neurol., vol. 55, no. 3, pp. 217–228, Mar. 2013, doi: 10.1111/j.1469-8749.2012.04436.x.

[35] R. H. Geuze, “Postural Control in Children With Developmental Coordination Disorder,”Neural Plast., vol. 12, no. 2–3, pp. 183–196, Jan. 2005, doi: 10.1155/NP.2005.183.

[36] “Geuze RH. Static balance and developmental coordination disorder. Hum Mov Sci. 2003 Nov;22(4-5):527–48. doi: 10.1016/j.humov.2003.09.008. PMID: 14624832.”.

[37] “Adams IL, Lust JM, Wilson PH, Steenbergen B. Compromised motor control in children with DCD: a deficit in the internal model?—A systematic review. Neurosci Biobehav Rev. 2014 Nov;47:225–44. doi: 10.1016/j.neubiorev.2014.08.011. Epub 2014 Sep 1. PMID: 25193246.”.

[38] “Kiemel T, Zhang Y, Jeka JJ. Identification of neural feedback for upright stance in humans: stabilization rather than sway minimization. J Neurosci. 2011 Oct 19;31(42):15144–53. doi: 10.1523/JNEUROSCI.1013-11.2011. PMID: 22016548; PMCID: PMC3470452.”

[39] “Bottaro A, Yasutake Y, Nomura T, Casadio M, Morasso P. Bounded stability of the quiet standing posture: an intermittent control model. Hum Mov Sci. 2008 Jun;27(3):473–95. doi: 10.1016/j.humov.2007.11.005. Epub 2008 Mar 14. PMID: 18342382.”.

[40] “Suzuki Y, Nomura T, Casadio M, Morasso P. Intermittent control with ankle, hip, and mixed strategies during quiet standing: a theoretical proposal based on a double inverted pendulum model. J Theor Biol. 2012 Oct 7;310:55–79. doi: 10.1016/j.jtbi.2012.06.019. Epub 2012 Jun 23. PMID: 22732276.”.

[41] “Mongold, Scott & Georgiev, Christian & Naeije, Gilles & Vander Ghinst, Marc & Stock, Matt & Bourguignon, Mathieu. (2024). Age-related changes in ultrasound-assessed muscle composition and postural stability. Scientific Reports. 14. 10.1038/s41598-024-69374-8.”.

[42] “Legrand T, Mongold SJ, Muller L, Naeije G, Ghinst MV, Bourguignon M. Cortical tracking of postural sways during standing balance. Sci Rep. 2024 Dec 3;14(1):30110. doi: 10.1038/s41598-024-81865-2. PMID: 39627308; PMCID: PMC11615285.”.

[43] E. Verbecque, L. Vereeck, and A. Hallemans, “Postural sway in children: A literature review,” Gait Posture, vol. 49, pp. 402–410, Sep. 2016, doi: 10.1016/j.gaitpost.2016.08.003.

[44] “Julienne A, Verbecque E, Besnard S. Normative data for instrumented posturography: a systematic review and meta-analysis. Front Hum Neurosci. 2024 Dec 18;18:1498107. doi: 10.3389/fnhum.2024.1498107. PMID: 39743990; PMCID: PMC11688309.”.

[45] M. L. Peterson, E. Christou, and K. S. Rosengren, “Children achieve adult-like sensory integration during stance at 12-years-old,” Gait Posture, vol. 23, no. 4, pp. 455–463, Jun. 2006, doi: 10.1016/j.gaitpost.2005.05.003.

[46] E. Verbecque, P. H. L. D. Costa, P. Meyns, K. Desloovere, L. Vereeck, and A. Hallemans, “Age-related changes in postural sway in preschoolers,” Gait Posture, vol. 44, pp. 116–122, Feb. 2016, doi: 10.1016/j.gaitpost.2015.11.016.

[47] “Rival C, Ceyte H, Olivier I. Developmental changes of static standing balance in children. Neurosci Lett. 2005 Mar 11;376(2):133–6. doi: 10.1016/j.neulet.2004.11.042. Epub 2004 Dec 9. PMID: 15698935.”.

[48] N. Kirshenbaum, C. Riach, and J. Starkes, “Non-linear development of postural control and strategy use in young children: a longitudinal study,” Exp. Brain Res., vol. 140, no. 4, pp. 420–431, Oct. 2001, doi: 10.1007/s002210100835.

[49] “García-Soidán JL, Leirós-Rodríguez R, Romo-Pérez V, García-Liñeira J. Accelerometric Assessment of Postural Balance in Children: A Systematic Review. Diagnostics (Basel). 2020 Dec 22;11(1):8. doi: 10.3390/diagnostics11010008. PMID: 33375206; PMCID: PMC7822105.”.

